# TREM2 limits necrotic core formation during atherogenesis by controlling macrophage survival and efferocytosis

**DOI:** 10.1101/2023.05.15.539977

**Authors:** Marie Piollet, Florentina Porsch, Giuseppe Rizzo, Frederieke Kapser, Dirk J.J. Schulz, Máté G. Kiss, Kai Schlepckow, Estrella Morenas-Rodriguez, Mustafa Orkun Sen, Julius Gropper, Melanie Roesch, Laura Göderle, Anastasiya Hladik, Sylvia Knapp, Marco Colonna, Rudolf Martini, Christian Haass, Alma Zernecke, Christoph J. Binder, Clément Cochain

**Author notes:** shared senior authorship; correspondence to: Alma Zernecke, Christoph J. Binder or Clément Cochain. shared first authorship (Marie Piollet, Florentina Porsch). shared second authorship (Giuseppe Rizzo, Frederieke Kapser, Dirk J.J. Schulz, Máté G. Kiss).

## Abstract

Atherosclerosis is a chronic disease of the vascular wall driven by lipid accumulation and inflammation in the intimal layer of arteries [1], [2], and its main complications, myocardial infarction and stroke, are the leading cause of mortality worldwide [3]. Recent studies have identified Triggering receptor expressed on myeloid cells 2 (TREM2), a lipid-sensing receptor regulating several key myeloid cell functions [4], as a highly expressed marker of macrophage foam cells in experimental and human atherosclerosis [5]. However, the function of TREM2 in the development of atherosclerosis is unknown. Here, we show that hematopoietic or global TREM2 deficiency increases necrotic core formation in early experimental atherosclerosis. We further demonstrate that TREM2 is essential for the efferocytosis capacities of macrophages, and to the survival of lipid-laden macrophages, altogether indicating a crucial role of TREM2 in maintaining the balance between foam cell death and their clearance in atherosclerotic lesions, thereby controlling plaque necrosis.

## Main

Atherosclerosis is a multifactorial, lipid-driven chronic inflammatory disease that with its associated complications, myocardial infarction and stroke, remains the leading cause of mortality globally [3]. It is initiated when lipoprotein particles, primarily low-density lipoproteins (LDL), accumulate in the intima of large and medium-sized arteries, where they can undergo modifications such as oxidation [6] or aggregation [2]. These processes render LDL proinflammatory and promote the recruitment of immune cells including monocytes, which differentiate into macrophages that perform a myriad of functions in the plaque. For example, they can take up modified lipoprotein particles, leading to the formation of foam cells, which are considered hallmark cells of atherosclerotic plaques [1], [2]. Additionally, macrophages play important roles in the clearance of cellular debris and apoptotic cells - a process called efferocytosis [1], [2]. Local death of foam cells combined with a progressive impairment of the efferocytotic capacity of macrophages promote the formation of necrotic areas in plaques, a feature of unstable lesions [7]. Recent single cell sequencing studies have revealed a significant degree of heterogeneity within the macrophage pool during atherosclerosis, including multiple distinct subsets [5]. However, the functional role of these distinct subsets, and whether they may differentially affect disease development, is currently unknown. These single cell RNA sequencing (scRNA-seq) studies of atherosclerotic plaques in mice [8], [9], [10], [11] and humans [12], [13] have revealed the existence of a subset of TREM2*-*expressing macrophages that show an enrichment for genes associated with pathways involved in lipid metabolism, tissue remodelling and oxidative stress while generally displaying a non-inflammatory gene expression signature. TREM2-expressing cells were subsequently found to correspond to foam cells [11]. Interestingly, TREM2-expressing macrophages with similar transcriptional profiles were found in multiple disease settings in mice and humans, including the infarcted myocardium [14], obesity [15], non-alcoholic fatty liver disease (NAFLD) [16], [17], Alzheimer’s disease [18], cancer [19] and others [4]. We have recently uncovered a protective role for TREM2-expressing macrophages in non-alcoholic steatohepatitis (NASH) [20]. However, the functional role of TREM2 in the development of atherosclerosis remains elusive. Here, we sought to address the function of TREM2 in experimental atherosclerosis, and in atherosclerosis-relevant macrophage functions.

We analyzed *Trem2* gene expression patterns in scRNA-seq data of total mouse aortic cells in *Ldlr*^*-/-*^ mice fed a normal chow or an atherogenic diet for 8, 16 or 26 weeks [21] (**Figure 1A-C**). High levels of *Trem2* were observed in macrophages in atherosclerotic conditions, consistent with previous observations [5], [8]. We also noted substantial expression of *Trem2* in a subset of phenotypically modulated smooth muscle cells (VSMC 2), albeit at a much lower level than in macrophages, and in scattered endothelial cells and fibroblasts (**Figure 1B-C**). In human atherosclerotic coronary arteries [22], *TREM2* expression was almost exclusively restricted to mononuclear phagocytes, but was also detected in some fibroblasts and fibromyocytes (**Figure 1D-F**). As we sought here to elucidate the role of TREM2 in bona fide macrophages from the hematopoietic compartment, and our previous data clearly support hematopoietic cells as a major source of TREM2 macrophages in dyslipidemia [20], we decided to employ *Ldlr*^*-/-*^ bone marrow chimeras with hematopoietic deficiency in *Trem2* as our main model (**Figure 2A**). Moreover, we used *Ldlr*^*-/-*^ rather than *Apoe*^*-/-*^ mice for atherogenesis experiments, as APOE is a downstream effector of TREM2 in phagocytes [23]. Briefly, lethally irradiated *Ldlr*^*-/-*^ mice had their hematopoietic compartment reconstituted with *Trem2*^*+/+*^ or *Trem2*^*-/-*^ bone marrow cells, and after a 4-6 week recovery period were fed an atherogenic diet (high fat diet: HFD; see **Methods**) for 8, 12, 16 or 20 weeks. These experiments were independently performed in Würzburg (Zernecke/Cochain labs, 8 and 20 weeks HFD) and in Vienna (Binder lab, 12 and 16 weeks HFD). After sacrifice, we examined atherosclerotic lesion formation and cellular plaque content in the aortic sinus as the primary readout site.

**Figure 1:**
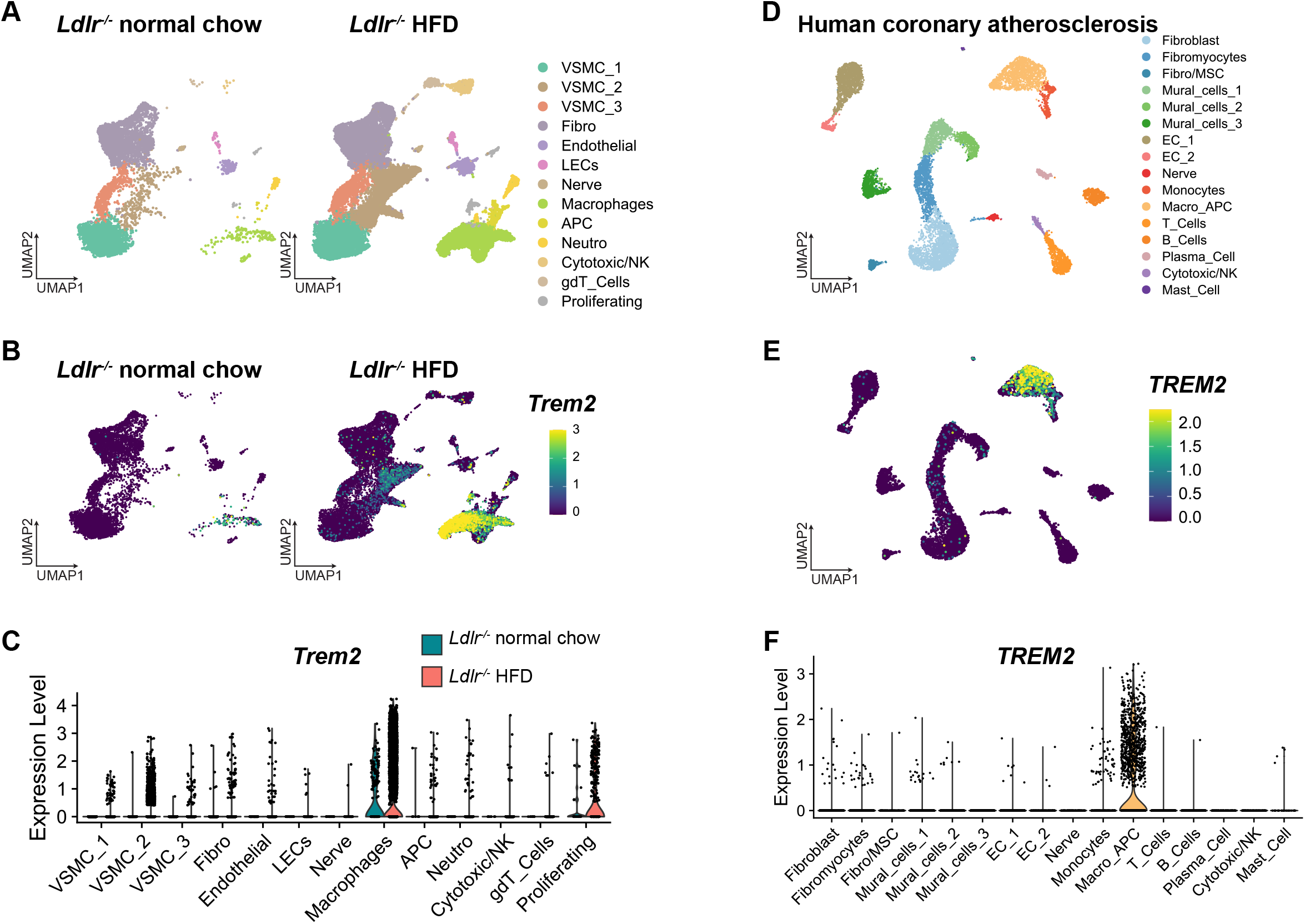
*Trem2* expression patterns at the single-cell level in mouse and human atherosclerosis. **A)** UMAP plot of total mouse aortic cells in *Ldlr*^*-/-*^ mice fed normal chow or a high fat diet (HFD) for 8, 16 or 26 weeks (*Ldlr*^*-/-*^ HFD); **B)** expression of *Trem2* in murine aortas projected onto the UMAP plot, split according to experimental condition; **C)** violin plot of *Trem2* expression in mouse aortic cells, split according to experimental conditions. **D)** UMAP plot of human atherosclerotic coronary artery cells (n=4 patients, data from [22]); **E)** expression of *TREM2* projected onto the UMAP plot and **F)** violin plot of *TREM2* expression across human atherosclerotic coronary artery cell populations. VSMC: vascular smooth muscle cells; Fibro: fibroblasts; LECs: lymphatic endothelial cells; APC: antigen presenting cells; Neutro: neutrophils; Cytotoxic/NK: cytotoxic T cells and natural killer cells; gdT: gammadelta T cells; MSC: mesenchymal stromal cells; EC: endothelial cells.

**Figure 2:**
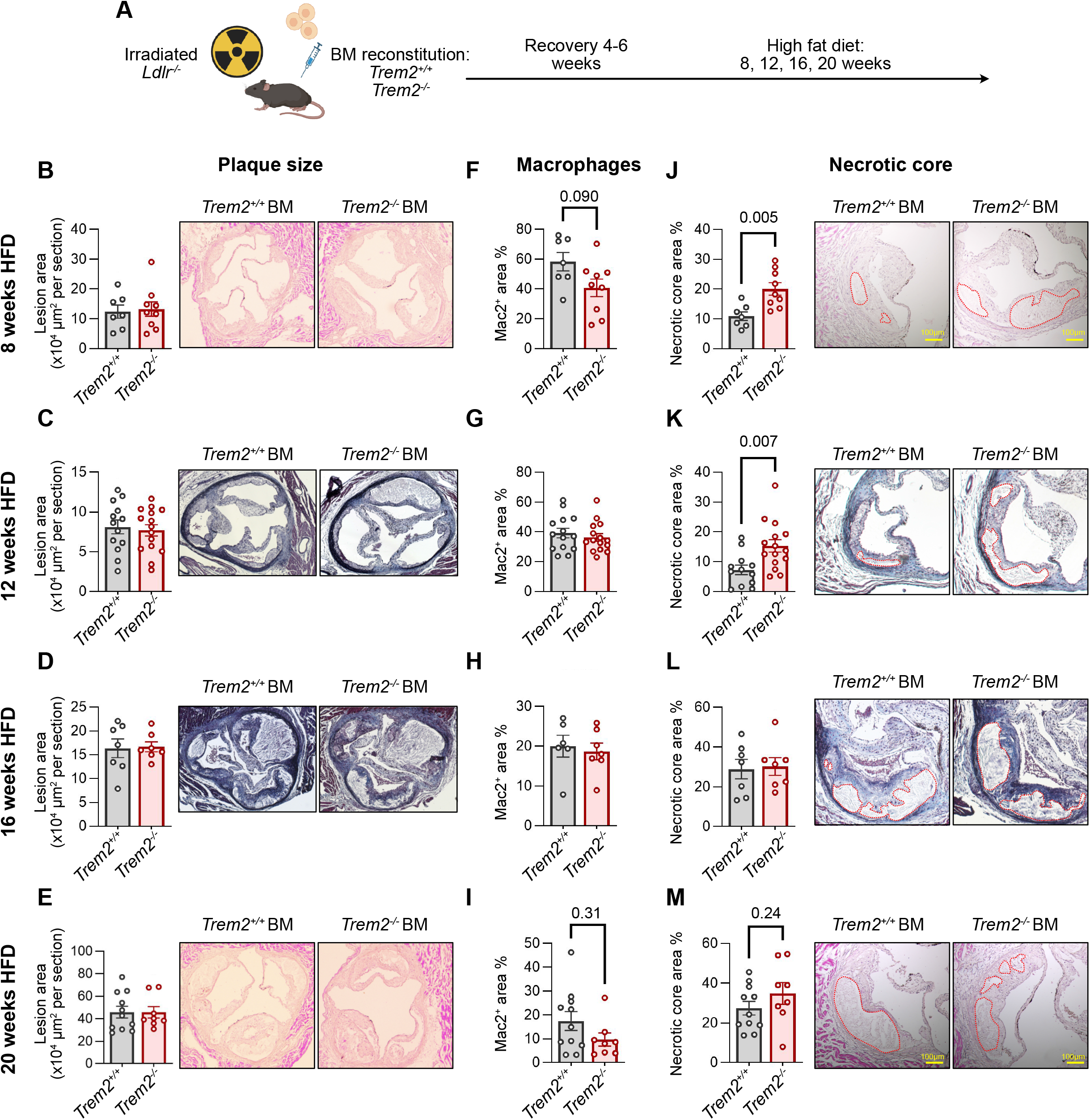
TREM2 controls necrotic core formation in early experimental atherosclerosis. **A)** experimental design for atherogenesis experiments in bone marrow chimeras reconstituted with *Trem2*^*+/+*^ or *Trem2*^*-/-*^ bone marrow. **B-M)** aortic sinus plaque size (**B-E**), macrophage content (expressed in percent of cellular plaque area; **F-I**), and necrotic core size (expressed in percent of total plaque area; necrotic area is depicted as a dashed red line; **J-M**) in *Ldlr*^*-/-*^ mice irradiated and reconstituted with *Trem2*^*+/+*^ or *Trem2*^*-/-*^ bone marrow cells and fed a high fat diet (HFD) for 8 weeks (**B, F, J**), 12 weeks (**C, G, K**), 16 weeks (**D, H, L**) or 20 weeks (**E, I, M**).

At all the time points analyzed, atherosclerotic lesion size in the aortic sinus was not affected by hematopoietic *Trem2* deficiency (**Figure 2B-E**). Additionally, we did not observe effects on lesion size at other vascular sites (aorta, innominate artery) except for an increased lesion size measured in *en face* aortas after 12 weeks of HFD in mice with hematopoietic *Trem2* deficiency (**Supplementary Figure 1A-B**). Mac2^+^ (i.e. *Lgals3*^*+*^) plaque macrophage coverage within the cellular areas of lesions was not affected, except at the earliest time point (8 weeks of HFD), where we noted a trend towards decreased macrophage content in mice transplanted with *Trem2*^*-/-*^ BM (p=0.09) (**Figure 2F-I**). During early lesion formation, we evidenced a strong increase in necrotic core size in mice with hematopoietic *Trem2* deficiency, with a 1.7-fold increase at 8 weeks (p<0.01) and a more than twofold increase at 12 weeks of HFD feeding (p<0.01) (**Figure 2J-K**). At 16 and 20 weeks of HFD, the necrotic core was expectedly larger than at earlier times, but no differences were observed across experimental groups (**Figure 2L-M**). While we chose BM chimeras as our primary model to avoid confounding effects of TREM2 expression on non-hematopoietic cells, we also analyzed atherogenesis in *Ldlr*^*-/-*^ *Trem2*^*-/-*^ mice after 10 and 20 weeks of HFD as a complementary model (**Supplementary Figure 2**). After 10 weeks of HFD, *Ldlr*^*-/-*^*Trem2*^*-/-*^ mice did not show any significant difference in plaque size in the aortic sinus and a trend towards increased lesion size in the aorta. Consistent with our BM chimera experiments, *Ldlr*^*-/-*^*Trem2*^*-/-*^ had a decreased macrophage content and an increased necrotic core area at this time point (**Supplementary Figure 2**). After 20 weeks of HFD, *Ldlr*^*-/-*^*Trem2*^*-/-*^ mice displayed a trend towards decreased plaque size and no differences in macrophage content or necrotic core formation (**Supplementary Figure 2**). Overall, data obtained in *Ldlr*^*-/-*^*Trem2*^*-/-*^ mice are in line with observations in BM chimera experiments, with an increased necrotic core size at early stages of lesion formation. Previous publications have suggested a role for TREM2 in systemic lipid metabolism and weight gain during diet-induced obesity, although conflicting evidence has been reported, suggesting either a protective [15], deleterious [24], or no influence [25] of TREM2 in this setting. In our BM chimera experiments and in *Ldlr*^*-/-*^*Trem2*^*-/-*^ mice, we did not observe significant effects on body weight at sacrifice, and inconsistent effects on systemic lipid levels, with *Trem2* deficiency being associated with no modification, slight increases, or slight reductions in total cholesterol and triglycerides in the serum across experiments (**Table S1**). Importantly, *Trem2* deficiency increased necrotic core formation in early atherosclerosis in mice with no changes in systemic lipid parameters (BM chimeras 12 weeks HFD and *Ldlr*^*-/-*^*Trem2*^*-/-*^ mice 10 weeks HFD), but also in mice showing decreased total cholesterol and triglyceride levels (*Trem2*^*-/-*^ BM chimeras, 8 weeks HFD), indicating that increased necrotic core formation is not driven by dyslipidemia. Altogether, these data indicate that TREM2 deficiency is associated with increased necrotic core formation during early experimental atherogenesis, independently of lesion size and systemic lipid levels.

Necrotic core formation is a hallmark of unstable atherosclerotic plaques, and results from a local imbalance between cell death, often associated with macrophage cholesterol overloading, and efferocytosis [7]. We tested whether TREM2 influences macrophage foam cell formation, their survival, and macrophage efferocytosis ability. Exposure of thioglycollate-elicited peritoneal macrophages to copper-oxidized LDL (Cu-OxLDL) induced a concentration-dependent expression of *Trem2* and other markers of the foamy/lipid-associated macrophage signature (*Gpnmb, Lgals3, Abca1* and *Abcg1*) (**Supplementary Figure 3A**). Macrophage uptake of Dil-labeled oxidized LDL (Dil-OxLDL) was reduced in *Trem2*^*-/-*^ bone marrow-derived macrophages (BMDM) (**Supplementary Figure 3B-C**), indicating a role of TREM2 in modified LDL uptake. We further evaluated macrophage survival in response to free cholesterol loading as an atherosclerosis-relevant *in vitro* model of macrophage cell death [26]. *Trem2*^*-/-*^ BMDM showed reduced survival in response to free cholesterol loading (**Figure 3A-C**), indicating that TREM2 promotes macrophage foam cell survival. In a flow cytometry-based efferocytosis assay, *Trem2*^*-/-*^ thioglycolate-elicited peritoneal macrophages showed a reduced ability to take up apoptotic cells (**Figure 3D-F**). Previous reports have shown that efferocytosis reprograms macrophage gene expression to promote continuous efferocytosis and resolution of inflammation, notably upregulating expression of the phagocytic receptor *Mertk* [27] and anti-inflammatory *Il10* [28]. Furthermore, the cellular lipid load induced by phagocytic challenge reprograms phagocytes to upregulate lipid metabolism-related genes in a TREM2-dependent manner [23]. While *Trem2*^*+/+*^ BMDM upregulated *Mertk, Il10*, the cholesterol efflux transporter *Abca1* and characteristic markers of the *Trem2*^*hi*^/foamy/lipid-associated macrophage signature (*Gpnmb, Fabp5*) [5] in response to apoptotic cell efferocytosis, this response was blunted in *Trem2*^*-/-*^ BMDM (**Figure 3G-H**). Based on our *in vivo* and *in vitro* observations, we propose the following model (**Figure 3I**): by enhancing oxLDL uptake and macrophage foam cell survival, and by increasing macrophage efferocytosis ability and anti-inflammatory gene expression in response to efferocytosis, TREM2 limits atherosclerotic plaque necrosis in early experimental atherogenesis.

**Figure 3:**
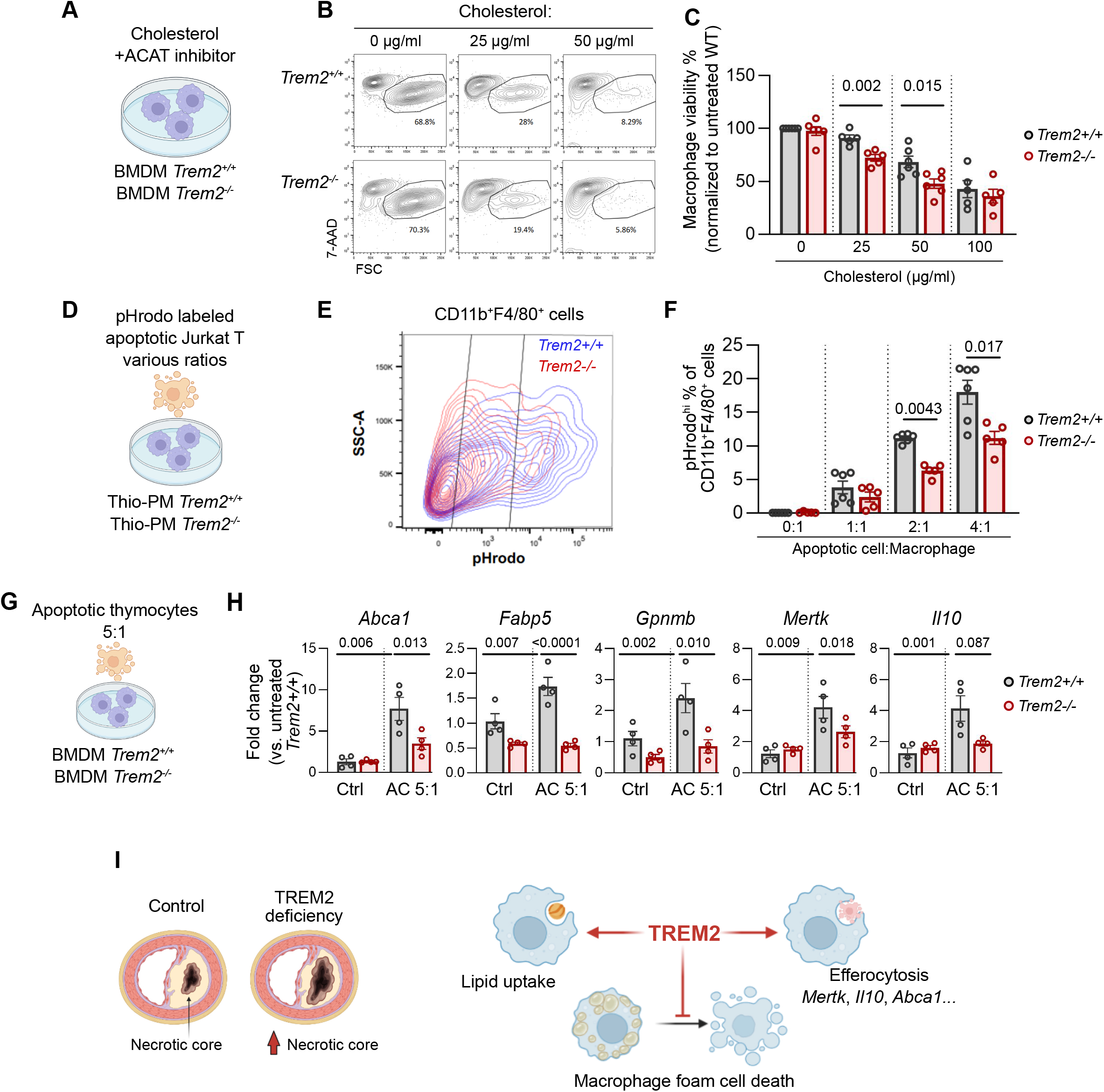
TREM2 controls macrophage survival and efferocytosis. **A)** Experimental design, **B)** representative flow cytometric analysis of bone marrow-derived macrophage (BMDM) survival in response to free cholesterol loading and **C)** analysis of macrophage viability (expressed as % of untreated WT control); each data point represents macrophages from one mouse assayed in technical triplicates; data were pooled from 5 experiments with 1 mouse from each genotype; **D-F)** in vitro efferocytosis assay with Thiogylcolate elicited peritoneal macrophages (Thio-PM) with **D)** experimental design, **E)** representative flow cytometry plot with a 4:1 apoptotic cells (AC):macrophages ratio, and **F)** quantitative analysis. **G-H)** gene expression in BMDM in response to efferocytosis with **G)** experimental setup and **H)** expression of the indicated transcripts expressed relative to untreated *Trem2*^*+/+*^ BMDMs; each data point represents macrophages from one mouse assayed in technical triplicates (AC 5:1: apoptotic thymocytes at a 5 thymocyte:1 macrophage ratio). **I)** Proposed model and overview of the conclusions.

A recent report investigating the role of TREM2 in macrophages in atherosclerosis proposed that TREM2 promotes lipid uptake and macrophage survival, and that TREM2 deficiency in macrophages decreases lesion formation *in vivo* [29], thereby showing TREM2-mediated modulation of macrophage functions consistent with our observations, but contrasting outcomes of *in vivo* disease readouts. Whether macrophage foam cell survival is deleterious or beneficial during atherogenesis likely depends on the disease stage [30], [31] and is difficult to investigate in experimental mouse models where complex lesions develop within short time frames (weeks) with continuous high levels of circulating LDL, compared to human lesions developing over decades [32]. In early atherosclerosis, the ability of macrophages to take up lipids and engage the appropriate mechanisms to clear them from the intima, while surviving and maintaining their efferocytosis ability, appears necessary to maintain local tissue homeostasis. By promoting the survival of lipid handling macrophages and macrophage efferocytosis, TREM2 would have a rather beneficial role in this context. Furthermore, while local macrophage death might limit plaque progression or underlie plaque regression upon reversal of dyslipidemia [33], this requires preserved efferocytosis and timely removal of dead cells. At later stages of atherosclerosis, other mechanisms might prevail in the regulation of macrophage accumulation, survival and efferocytosis, with macrophage proliferation (also potentially regulated by TREM2 [29]) becoming a major pathway of macrophage accumulation [34], and inhibition of Mertk-dependent efferocytosis underlying plaque necrosis [35]. Our observations that TREM2 modulates macrophage survival and efferocytosis are overall consistent with previously proposed roles of TREM2 in other pathological contexts. Indeed, TREM2, a marker of disease associated microglia in the brain [18], has been proposed to promote survival of these cells in conditions of metabolic stress [36], promotes efferocytosis in the diseased liver [37] and after ischemic stroke [38], and regulates expression of genes involved in lipid handling following a phagocytic challenge [23]. Another recent report also proposed that TREM2 promotes macrophage lipid uptake, consistent with our results, but showed reduced atherogenesis in *Apoe*^*-/-*^*Trem2*^*-/-*^ mice [39]. These contrasting findings might be caused by the use of *Apoe*^*-/-*^ mice, as APOE is a known downstream effector of TREM2 in phagocytes [23], and macrophage-derived APOE controls macrophage survival, apoptotic cell clearance and necrotic core formation in atherosclerosis [31].

Besides its local role in lesional macrophages, previous evidence indicates that TREM2 functions in other organs might in addition impact on atherosclerosis. In diet-induced obesity, TREM2 has been proposed to control adipose tissue remodeling [25] and possibly affect metabolic homeostasis systemically [15]. Furthermore, TREM2 has a protective role in the liver in experimental NASH [20], [37]. We did not observe consistent effects of TREM2 deficiency on weight gain or systemic lipid levels under atherogenic conditions, and we observed an increased necrotic core formation in mice with similar or even decreased systemic lipid levels, indicating that TREM2 limits plaque necrosis independently of lipid levels. However, we cannot fully exclude that TREM2 functions in other organs might have influenced lesion formation in Trem2-/-mice, as for instance brain microglia activity, which is highly TREM2-dependent, might be involved in atherosclerosis [40]. While *Trem2* expression is mostly restricted to mononuclear phagocytes in health and disease [4], we observed expression of *Trem2* in phenotypically modulated VSMCs in scRNA-seq. Guo et al. proposed that TREM2 promotes lipid uptake by VSMCs [39]. Uncovering a potential role of TREM2 in phenotypically modulated VSMCs during atherogenesis, however, requires further studies using appropriate cell-type specific models.

In conclusion, our data uncover a role of TREM2 as a regulator of macrophage survival and efferocytosis in atherosclerosis, thereby limiting plaque necrosis. TREM2 might thus represent an attractive therapeutic target in cardiometabolic disease, as in addition to its proposed beneficial roles in obesity and other lipid-driven diseases such as NASH, TREM2 may limit necrotic core formation in atherosclerotic lesions.

## Acknoledgements

This work was supported by the Interdisciplinary Center for Clinical Research (Interdisziplinäres Zentrum für Klinische Forschung (IZKF), University Hospital Würzburg (A-384 to A.Z.), the Deutsche Forschungsgemeinschaft (DFG; German Research Foundation, Projects 374031971-TRR 240, 324392634-TR221, ZE827/14-1, project numbers 396923792 to A.Z., 432915089 to A.Z. and C.C., 458539578 and 471705758 to C.C.), the DFG SFB1525 project number 453989101 (projects A1 and B3 to A.Z., project A6 and PS2 to C.C.). This work was supported by the Deutsche Forschungsgemeinschaft (DFG, German Research Foundation) under Germany’s Excellence Strategy within the framework of the Munich Cluster for Systems Neurology (EXC 2145 SyNergy – ID 390857198) and a Koselleck Project HA1737/16-1 (to CH).

