## Supplemental_Methods_Figures for "TREM2 limits necrotic core formation during atherogenesis by controlling macrophage survival and efferocytosis"

**Supplementary Information**

- Supplementary Online Methods
- Supplementary Table S1
- Supplementary Figure 1-3

**Materials and Methods**

**Animal models**

Mice: *Ldlr*$^{-/-}$ mice (B6.129S7-Ldlrtm1Her/J, JAX stock no. 002207) were obtained from the Jackson Laboratory (Bar Harbor, USA). *Trem2*$^{-/-}$ mice were provided by Marco Colonna (Washington University, St. Louis, USA). *Trem2*$^{-/-}$ *Ldlr*$^{-/-}$ mice were obtained by crossing the above mouse strains in-house. All mice were on a C57BL/6J background.

Würzburg experiments: Mice were bred and kept in individually-ventilated cages (IVC) with a 12-hour dark-/light-cycle and ad libitum access to sterilized food and water under barrier-specific pathogen-free conditions (SPF). Six to eight-week-old male or female *Ldlr*$^{-/-}$ and *Ldlr*$^{-/-}$*Trem2*$^{-/-}$ mice were fed with an atherogenic diet (15% milk fat, 1.25% cholesterol; Altromin) for 10 or 20 weeks. Bone marrow chimeras were generated by lethally irradiating 6 to 8 week old male *Ldlr*$^{-/-}$ mice (9 Gy, Faxitron CP-160). 4 hours after irradiation, the mice received 5x10$^{6}$ total bone marrow cells from *Trem2*$^{+/+}$ or *Trem2*$^{-/-}$ donors, and were left to recover for 4 weeks, with neomycin sulfate (bela-pharm, 2 g/l in drinking water) as antibiotic prophylaxis during the first week. Afterwards, mice were fed with an atherogenic diet (15% milk fat, 1.25% cholesterol; Altromin) for 8 or 20 weeks. All animal studies conform to the Directive 2010/63/EU of the European Parliament and have been approved by the appropriate local authorities (Regierung von Unterfranken, Wuerzburg, Germany, Akt.-Z. 55.2-2531.01-24/13, 55.2-DMS-2532-2-287, 55.2-DMS-2532-2-1227).

Vienna Experiments: Age-matched male mice of at least 8 weeks of age were used for all experiments and groups were kept co-housed. Mice were bred and kept in individually ventilated cages (IVC) with a 12-hour dark-/light-cycle and *ad libitum* access to sterilized food and water under barrier-specific pathogen-free conditions (SPF) at the Core Facility for Animal Breeding and Husbandry of the Center for Biomedical Research at the Medical University of Vienna, Austria. For bone marrow transplantation studies, 8 week old *Ldlr*$^{-/-}$ mice were **γ**-irradiated using two doses of 6 Gy spaced 4 hours apart to eliminate haematopoietic cells. The bone marrow was subsequently reconstituted with 5x10$^{6}$ bone marrow cells isolated from the femurs and tibias of 6-8 week old *Trem2*$^{-/-}$ donor mice or *Trem2*$^{+/+}$ littermates. Recipient mice recovered for 6 weeks post bone marrow transplantation to allow for reconstitution of the haematopoietic compartment. To induce atherosclerosis, mice were put on a high-fat high-cholesterol diet containing 21% milk fat and 0.21% cholesterol (TD88137, Ssniff Spezialdiäten GmbH, Soest, Germany) with *ad libitum* access to pellets for 12 or 16 weeks. The diet pellets were previously irradiated/autoclaved for sterility. All experimental studies were approved by the Animal Ethics Committee of the Medical University of Vienna and the Austrian Federal Ministry of Education, Science and Research, and were performed according to Good Scientific Practice and national and international institutional guidelines (License number BMWF 66.009/0336-V/3b/2018).

**Histology**

<u>*Würzburg experiments:*</u>

*Aorta Oil-Red-O staining for atherosclerotic lesion quantification*

Mice were killed by cervical dislocation under Isoflurane anesthesia. Aorta was perfused with PBS, excised and fixed in 4% Paraformaldehyde (PFA) for 24 hours. After removing the adventitia layer, the aortas were washed in PBS for 5 minutes and dipped 10 times in 60% 2-propanol. Oil-Red-O staining was performed incubating the aortas for 15 minutes in Oil-Red-O staining solution. After washing with 60% 2-propanol and PBS, the aortas were mounted and pictures were acquired with Leica DM 4000 B LED microscope. Lesion size was assessed measuring the red staining area using ImageJ software (Fiji).

*Aortic root preparation and histology staining*

Mice were killed by cervical dislocation under Isoflurane anesthesia. The heart was exposed, perfused with PBS, excised and fixed in 4% PFA for 24 hours. Aortic root sections were made with Cryostat (Leica, CM3050 S) at 4 µm thickness. For immunofluorescence staining, antigen retrieval with citrate method was performed. The sections were then blocked for 30 minutes with blocking solution containing 2% mouse serum, 2% rabbit serum, 2% horse serum, 1% BSA, and 0,1% tritonX-100 to prevent unspecific staining. Then, they were incubated with rat anti-mouse MAC2 (Cedralane, CL8942AP, M3/38) antibodies overnight at 4°C. After washing with PBS, sections were stained with goat anti-rat AlexaFluor488 (ThermoFisher, A1106). Finally, the sections were mounted using Vectashield (Vector Laboratories, H1200) containing DAPI and pictures were acquired with Leica DM 4000 B LED microscope. MAC2 fluorescence area was measured using ImageJ software. For necrotic core measurement, hematoxylin and eosin staining was performed on aortic root sections. Briefly, the sections were stained with hematoxylin (Morphisto, 10231) solution for 6 minutes, washed shortly in distilled water and then in running tap water for 6 minutes. Aortic root sections were stained with eosin (Morphhisto, 10177) for 6 minutes and then washed shortly in distilled water. The sections were dehydrated in increased ethanol concentration and xylene as the last step. The slides were mounted with a mounting medium and pictures were acquired with Leica DM 4000 B LED microscope. Necrotic core area and plaque size were measured using ImageJ software.

*Vienna experiments:*

*Assessment of atherosclerosis*

For *en face* analysis, aortas ranging from the aortic arch to the iliac bifurcation were isolated from mice post mortem and stained with Sudan IV (Sigma-Aldrich, St. Louis, USA). Lesion size was assessed in a double-blinded fashion by computer-assisted analysis using Photoshop Elements (Adobe Inc, San Jose, USA) and expressed as % of Sudan IV+ area per total aortic area. For aortic root cross sections, hearts were isolated post mortem, fixed and embedded in paraffin. Aortic roots were sectioned in 5 µm-thick serial sections starting from the appearance of the three aortic valve leaflets. 9 sections separated by 50 µm across a distance of 400 µm from aortic root origin were used for subsequent analysis. Sections were stained using Masson Trichrome staining according to manufacturer's instructions (Sigma-Aldrich, St. Louis, USA) and photographed on a Zeiss AxioImager A1 using AxioCam MRC5 and the Zen2.3 Pro Softwarte (Carl Zeiss AG, Jena, Germany). Lesion size in all three leaflets was assessed in a double-blinded fashion by computer-assisted image analysis using Photoshop Elements (Adobe Inc, San Jose, USA) and ImageJ software. For necrotic cores, data are expressed as % of lesional area made up of necrotic, acellular area. Necrotic cores were assessed at 150-250 µm from aortic root origin. For innominate arteries, brachiocephalic arteries were isolated from the mice post mortem, fixed and embedded in paraffin and sectioned in 5 µm-thick serial sections. 5 sections separated by 50 µm were used for subsequent analysis in a similar fashion as aortic root cross sections. Lesion size is expressed as % stenosis (i.e. % of artery luminal surface covered by atherosclerotic lesions).

*Immunohistochemistry*

For the assessment of lesional macrophage content, aortic root cross sections were stained with anti-Mac2 antibody (BioLegend, San Diego, USA). Quantification was performed by image-assisted analysis using ImageJ software to determine Sirius-Red-positive or Mac2-positive areas, respectively. Data are expressed as % of positive areas within the cellular areas of the atherosclerotic plaques.

**Blood cholesterol and triglyceride measurements**

*Würzburg experiments:*

Blood was collected in serum collection tubes (Starsted, 41-1500-005) and kept on ice until all samples were collected. After equilibrating samples to room temperature for 30 minutes, they were centrifuged at 10,000 rcf for 5 minutes. The serum was aliquoted and stored at -80°C until further use. Total cholesterol was measured using Amplex Red Cholesterol Assay kit (Invitrogen, A12216) according to manufacturer's instructions. Optical density was measured using Microplate Reader. Triglycerides were measured using EzymChrom$^{TM}$ Triglyceride Assay kit (BioAssaysSystems, ETGA-200) according to manufacturer's instructions. Optical density was measured using Microplate Reader.

*Vienna experiments:*

Blood was collected using 23-gauge needles post mortem after >4 hours of fasting via the vena cava into EDTA collection tubes (Greiner Bio-One, Germany). Plasma was obtained by centrifugation at 1000xg for 20 minutes at room temperature. Plasma triglyceride and cholesterol levels were measured in an ISO-15189 accredited medical laboratory under standardized conditions on Beckman Coulter AU5400 instruments using Beckman Coulter OSR6516 Reagent (Beckman Coulter, Brea, USA) at the Department of Laboratory Medicine, Medical University Vienna, Austria. Alternatively, plasma lipid content was measured according to manufacturer's instructions using Liquid Reagents kit (GPO-PAP Triglyceride Liquicolor kit, CHOD-PAP Cholesterol Liquicolor kit, HUMAN Biochemica and Diagnostica mbH, Wiesbaden, Germany).

**Bone marrow-derived macrophages**

Femur and tibia of two hindlimbs were used for bone marrow isolation as described in [1]. Bone marrow cells were resuspended in RPMI supplemented with 10% FCS, 100 U/ml penicillin/streptomycin, and 50 µM β-Mercaptoethanol), filtered (70 µm cell strainer) and washed. Cells were resuspended in medium supplemented with 15% of L929 conditioned medium, cells were counted, and $2x10^6$/ml cells were plated in a $10cm^2$ cell culture dish. After 7 days, macrophages were detached using Accutase (Sigma-Aldrich, A6964), washed, resuspended in 15% L929 supplemented medium, and $0.4x10^6$ macrophages were plated per well in a 12-well plate. Cells were rested overnight before conducting experiments. Before all in vitro assays, macrophages were incubated for 4 hours in starving low-serum medium (RPMI supplemented with 0.5% FCS)

**Thioglycoloate-elicited Peritoneal Macrophages**

To elicit peritoneal macrophages, 12-16 week old male mice received a single dose (50ul/ug body weight) of sterile thioglycolate (Thermo Fisher Scientific, Difco Laboratories, Waltham, MA, USA) by intraperitoneal injection 72 hours prior to sacrifice. After sacrifice, thioglycolate-elicited macrophages were harvested by peritoneal lavage using sterile PBS+1%BSA. Cells were subsequently plated in RPMI-1640 medium supplemented with 10% heat-inactivated FCS and were allowed to adhere for 2 hours, after which cells were used for subsequent experiments (oxLDL loading, efferocytosis assays).

**Ox-LDL uptake assay**

*Würzburg experiments:*

Macrophages were exposed to 5 µg/ml of Dil-Ox-LDL (Thermofisher, K1612) overnight. Cells were washed once with PBS and detached with Accutase. Macrophages were then collected in FACS tube, washed with PBS supplemented with 1% FCS, and incubated in Fc Block (1:50, Biolegend, 101320) for 10 minutes. After blocking, macrophages were stained with F4/80 e450 (1:100, eBioscience, 48-4801-82, BM8) and Viability dye e780 (1:1000, thermofisher, 65-0865-14) for 30 minutes in the dark. After washing, they were resuspended in PBS supplemented with 1% FCS and read by FACS Celesta (BD) and analyzed with Flowjo v10.

*Vienna experiments:*

Foam cell formation assays were performed as previously described [2]. Briefly, thioglycolate-elicited peritoneal macrophages were treated with 50µg/ml of Cu-OxLDL or 50µg/ml native LDL for 24 hours. Cells were subsequently rinsed in serum-free PBS and subsequently processed for RNA isolation.

**Viability assay**

Adherent macrophages were incubated with ACAT inhibitor (Sigma-Aldrich, S9318) at 2 µg/ml, together with different concentrations of soluble cholesterol (Sigma Aldrich, C4951) (25, 50 and 100 µg/ml). After 24h, the cells were washed with PBS then detached gently with accutase and stained with 7AAD (Thermofisher), according to the manufacturer's instructions. The cells were read by FACS Celesta (BD) and analyzed with Flowjo v10.

**RNA Isolation, cDNA generation and assessment of gene expression in response to cholesterol loading** (Vienna)

For RNA extraction from in vitro OxLDL-treated macrophages, cells were lysed according to manufacturer's instructions using QIAzol Lysis Reagent (Qiagen, Hilden, Germany). In order to generate cDNA for subsequent quantitative real-time PCR experiments, up to 0,3 µg of RNA was subsequently reverse transcribed according to manufacturer's instructions using the High-Capacity cDNA Reverse Transcription kit (Applied Biosystems, Thermo Fisher, Waltham, MA, USA). Real-time PCR was subsequently performed on a CFX96 Real Time

PCR System (BioRad Laboratories, Hercules, CA, USA) using a KAPA SYBR Fast kit (Thermo Fisher) and the primers indicated in the table below. Gene expression was normalized to 18S.

| **Gene (mouse)** | **Forward Sequence (5'-3')** | **Reverse Sequence (3'-5')** |
|---|---|---|
| *18S* | AGTCCCTGCCCTTTGTACACA | CGATCCCAGGGCCTCACTA |
| *Trem2* | CTACCAGTGTCAGAGTCTCCGA | CCTCGAAACTCGATGACTCCTC |
| *Gpnmb* | GGCTACTTCAGAGCCACCATCA | CTTTGCAGGTCACAGTGAAGTCC |
| *Lgals3* | AACACGAAGCAGGACAATAACTGG | GCAGTAGGTGAGCATCGTTGAC |
| *Cd36* | GCCAAGCTATTGCGACATGA | AAAAGAATCTCAATGTCCGAGACTTT |
| *Abca1* | GGAGCCTTTGTGGAACTCTTCC | CGCTCTCTTCAGCCACTTTGAG |
| *Abcg1* | GACACCGATGTGAACCCGTTTC | GCATGATGCTGAGGAAGGTCCT |

**Efferocytosis Assay** (Vienna)

To generate apoptotic bait cells, Jurkat cells were rendered apoptotic by exposure to UV-irradiation (100mJ/cm2) and kept in culture at 37°C with 5% CO2 for 24 hours prior to the efferocytosis assay. Jurkat cells were subsequently stained using pH-sensitive pHrodo Red dye (Thermo Fisher) and cultured with previously plated adherent thioglycolate-elicited peritoneal macrophages at the indicated ratios (1:1, 2:1 and 4:1). pHrodo-labelled apoptotic cells were incubated with macrophages for 2 hours, after which cells were removed and remaining unbound apoptotic cells were washed off using PBS. Macrophages were subsequently harvested and stained (anti-CD11b, anti-F4/80) in flow-cytometry buffer (cold PBS supplemented with 1% BSA) to distinguish macrophages from remaining bait cells. Macrophages were assessed for apoptotic cell uptake by flow cytometry-based measurement of pHrodo-positivity of CD11b+ F4/80+ cells.

**Gene expression in response to efferocytosis**

Cultured macrophages were incubated for 16 hours either with apoptotic thymocytes at a 5 thymocyte:1 macrophage ratio or left untreated. Thymocytes had been previously extracted from the thymus of C57BL6/J mice and rendered apoptotic by overnight incubation at 37°C in

1 µM of Staurosporine in RPMI supplemented with 10% FCS. After extensive washing to remove unbound thymocytes, adherent macrophages were lyzed in RA1 lysis buffer (with added β-mercaptoethanol) from the NucleoSpin RNA extraction Kit (Macherey-Nagel, 740855.50). Total RNA was extracted using the NucleoSpin RNA extraction Kit in accordance with the manufacturer's instructions. Equal amounts of template RNA were used for cDNA synthesis, and RNA was reverse transcribed using random hexamer primers of the First Strand cDNA Synthesis kit (Thermofisher, K1612). Quantitative real-time polymerase chain reaction was performed on triplicate samples of template cDNA with PowerUp™ SYBR™ Green Master Mix (Applied Biosystems, A25742) on an Applied Biosystems™ QuantStudio™ 6 Flex Real-Time PCR System using specific primer pairs. Quantitative measurements were determined using the ΔΔCt method, with *Hprt* as the housekeeping gene. Primer sequences were as follows:

| ***Gene*** | **Forward (5'->3')** | **Reverse (5'->3')** |
|---|---|---|
| *Hprt* | TCCTCCTCAGACCGCTTTT | CCTGGTTCATCATCGCTAATC |
| *Fabp5* | AAGCCACGGCTTTGAGGAGT | TTCACTGTGCTCTCGGTTTTG |
| *Gpnmb* | GGGCCATGAACAGTATCCCG | CCTTCTGGCATCTGGGGAAC |
| *Abca1* | AGTGATAATCAAAGTCAAAGGGACAC | AGCAACTTGGCACTAGTAACTCTG |
| *Il10* | ATTTGAATTCCCTGGGTGAGAAG | CACAGGGGAGAAATCGATGACA |
| *Mertk* | CAGGGCCTTTACCAGGGAGA | TGTGTGCTGGATGTGATCTTC |

**Single-cell RNA-seq analysis**

*Mouse scRNA-seq data:* sequencing data from Pan et al. Circulation 2020 [3] were downloaded from Gene Expression Omnibus GSE155513, pre-processed in cellranger-6.1.2, and further analyzed in Seurat v4.3.0 [4]. We used data from *Ldlr*$^{-/-}$ mice fed normal chow or a western diet for 8, 16 or 26 weeks (i.e. the following data from Gene Expression Omnibus

GSE155513: GSM4705592, GSM4705593, GSM4705594, GSM4705595, GSM4705596, GSM4705597, GSM4705598, GSM4705599). Individual datasets were pre-processed with quality control filtering in Seurat: cells containing >200 detected genes, and genes detected in at least 3 cells were included in the analysis using the 'CreateSeuratObject' function with 'min.features = 200' and 'min.cells=3'. Quality control filtering was further performed to remove dead/damaged cells with a high proportion of mitochondrial transcripts (>10%), and outlier cells with high UMI numbers. All data were log normalized using the 'NormalizeData' function in Seurat with default parameters. Data were pooled and batch corrected using Harmony [5] within Seurat. 2,000 highly variable genes were identified using 'FindVariableFeatures' (with selection.method = "vst"). Data were scaled using 'ScaleData' with default parameters, and principal component analysis performed using 'RunPCA' with default parameters, and batch corrected using 'RunHarmony' with default parameters. Dimensional reduction was performed using 'RunUMAP(reduction = "harmony", dims = 1:20)', and clustering was performed at a 0.4 resolution using 'FindNeighbors(reduction = "harmony", dims = 1:20)' followed by 'FindClusters(resolution = 0.4)'. Positive marker genes for each cluster were identified using 'FindAllMarkers'.

*Human scRNA-seq data*: data from total cells of human atherosclerotic coronary arteries [6] were analyzed in Seurat v3 [7] starting from the author provided cell-count matrix (downloaded from Gene Expression Omnibus GSE131778). Cells containing <200 detected genes were excluded, and genes detected in at least 3 cells were included in the analysis using the 'CreateSeuratObject' functions with 'min.features = 200' and 'min.cells=3'. Further quality control filtering was performed and cells with >5% mitochondrial transcripts were excluded, as well as cells with outlier number of UMIs (nCount_RNA>15,000). A total of 10,934 cells were analyzed. As a pre-analysis indicated a substantial patient-driven batch effect, we performed batch correction using Harmony [5] within Seurat, considering each patient as an independent sample. Data were normalized using the 'NormalizeData' function in Seurat with default parameters. 2,000 highly variable genes were identified using 'FindVariableFeatures' (with selection.method = "vst"). Data were scaled using 'ScaleData' with default parameters, and principal component analysis performed using 'RunPCA' with default parameters, and batch corrected using 'RunHarmony' with default parameters. Dimensional reduction was performed using 'RunUMAP(reduction = "harmony", dims = 1:20)', and clustering was performed at a 0.4 resolution using 'FindNeighbors(reduction = "harmony", dims = 1:20)' followed by 'FindClusters(resolution = 0.4)'. Positive marker genes for each cluster were identified using 'FindAllMarkers'. Cell type annotation was performed based on expression of known cell lineage markers, and on cluster annotations in [6].

### Statistical analysis

Statistical analyses were performed using GraphPad Prism version 9. Results are expressed as mean ± standard error (SEM). For two-group comparisons, normal distribution of the data was assessed by D'Agostino-Pearson test followed by an unpaired t-test (normally distributed data) or a non-parametric Mann-Witney test (non-normally distributed data). Data with multiple comparisons were assessed by one-way ANOVA followed by Tukey's test for multiple comparisons. P values $<0.05$ were considered statistically significant.

| | | Body weight mean (g) | | Total cholesterol (mg/dL) | | Triglycerides (mg/dL) | |
|---|---|---|---|---|---|---|---|
| Experimental Design | time HFD | *Trem2+/+ vs. Trem2-/-* | | *Trem2+/+ vs. Trem2-/-* | | *Trem2+/+ vs. Trem2-/-* | |
| BM Chimeras | 8 weeks | 26.0+/- 0.8 vs. 26.1+/-0.6 | ns | 1336.8+/-75.9 vs. 1114.6+/-51.5 | * | 405.3+/-37.9 vs. 309.2+/-17.6 | * |
| BM Chimeras | 12 weeks | 29.8+/-0.8 vs. 28.4+/-0.5 | ns | 1028.6+/-95.9 vs. 960.8+/-70.7 | ns | 556.7+/-57.8 vs. 554.5+/-48.8 | ns |
| BM Chimeras | 16 weeks | 29.7+/-1.2 vs. 29.5+/-0.8 | ns | 1337.5+/-107.9 vs. 1470.0+/-43.8 | ns | 707.1+/-40.3 vs. 780.4+/-64.0 | ns |
| BM Chimeras | 20 weeks | 24.1+/-0.4 vs. 24.9+/-0.5 | ns | 1558.8+/-59.0 vs. 1851.3+/-87.2 | * | data not available | N/A |
| *Ldlr-/- Trem2-/-* | 10 weeks | 31.1+/-1.7 vs. 29.9+/-1.4 | ns | 1730.7+/-78.0 vs. 1740.0+/-79.4 | ns | 507.6+/-91.1 vs. 532.2+/-98.1 | ns |
| *Ldlr-/- Trem2-/-* | 20 weeks | 33.9+/-1.5 vs. 32.0+/-1.2 | ns | 1302.9+/-101.5 vs. 1582.8+/-105.5 | ns | data not available | N/A |

all data: mean +/-SEM; *p<0.05 (Mann-Whitney U test); ns: not significant; N/A not available

### Table S1

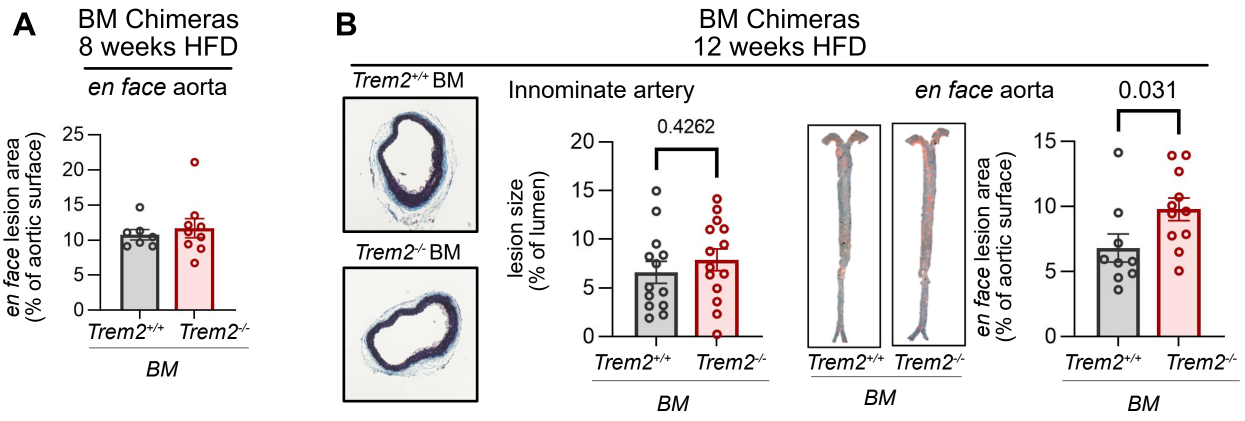


**Supplementary Figure 1: Analysis of additional vascular sites in bone marrow chimeras. A)** atherosclerotic lesion formation in the aorta at 8 weeks of HFD feeding and **B)** atherosclerotic lesion formation in the innominate artery and aorta at 12 weeks of HFD in *Ldlr*$^{-/-}$ mice irradiated and reconstituted with *Trem2*$^{+/+}$ or *Trem2*$^{-/-}$ bone marrow.

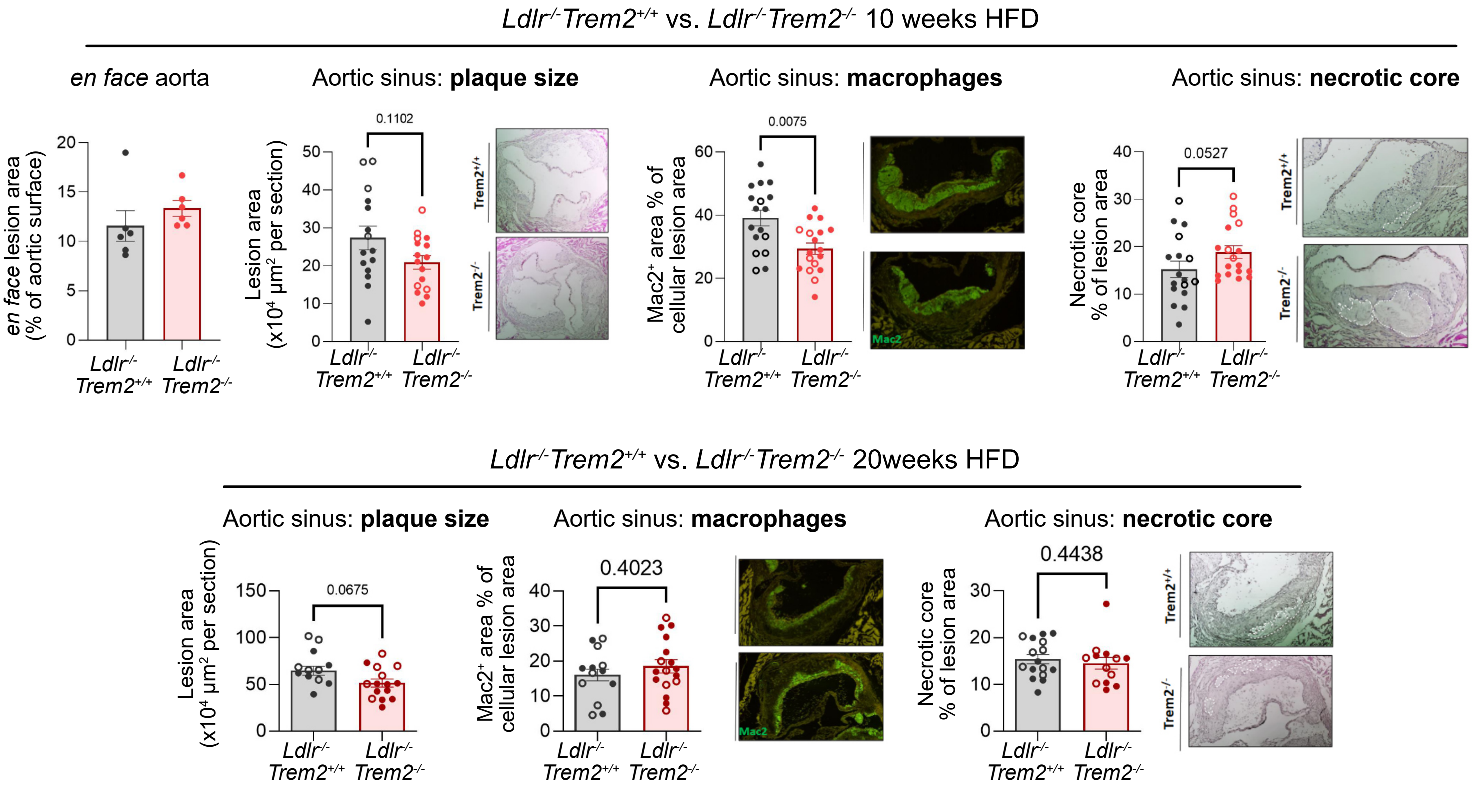


**Supplementary Figure 2: Analysis of atherosclerosis in *Ldlr*$^{-/-}$*Trem2*$^{-/-}$ mice.** Analysis of atherosclerosis in *Ldlr*$^{-/-}$*Trem2*$^{-/-}$ mice after 10 weeks (top) or 20 weeks (bottom) of HFD feeding. Open circles: female mice; filled circles: male mice

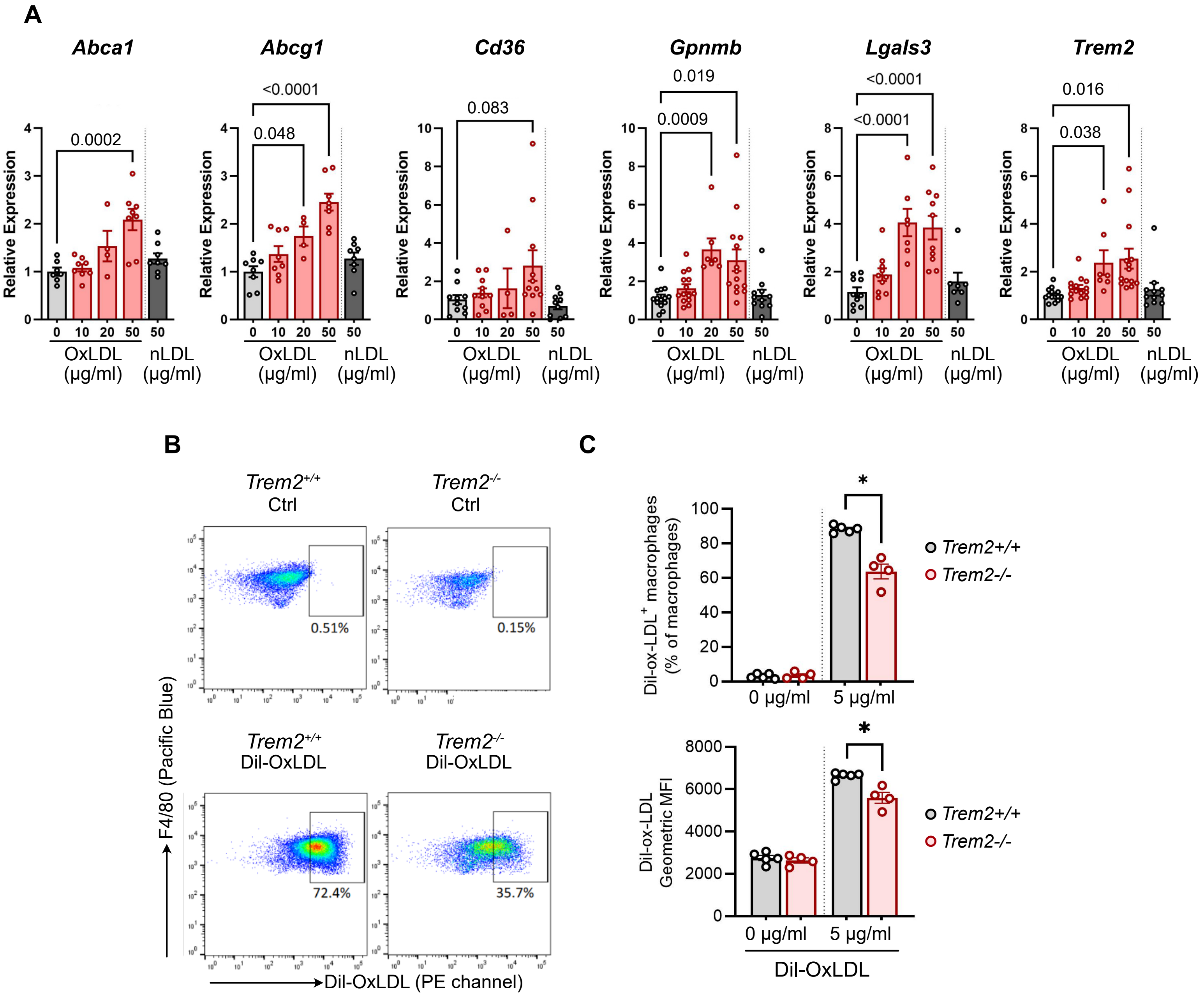


**Supplemental Figure 3: A)** Gene expression in thioglycollate elicited peritoneal macrophages in response to OxLDL and native LDL (nLDL); each data point represents macrophages from 1 mouse, pooled from 2-4 experiments; **B-C)** Dil-OxLDL uptake by bone marrow-derived macrophages assayed by flow cytometry. Each data point represents data from one mouse; representative of 3 independent experiments.
